# A small number of human lineage mutations regulated RNA-protein binding of conserved genes and promoted human evolution

**DOI:** 10.1101/2023.03.27.534315

**Authors:** Weichen Song, Shunying Yu, Min Zhao, Guan Ning Lin

## Abstract

Whether human lineage mutations (HLM) have contributed to human evolution via post-transcriptional modification remained unknown. We applied deep learning models Seqweaver to predict how HLM impacts RNA-binding protein affinity (RBP). At the threshold of the top 1% of human common variation, only 0.27% of HLM had large impacts on RBP. These HLMs enriched in a set of conserved genes that are highly expressed in adult excitatory neurons and prenatal Purkinje neurons, and involved in synapse organization and the GTPase pathway. These genes also carried excess damaging coding mutations that caused neurodevelopmental disorders, ataxia, and Schizophrenia. Among these genes, NTRK2 and ITPR1 had the most aggregated evidence of functional importance, pointing to an essential role in cognition and bipedalism. We concluded that a very small number of human-specific mutations have contributed to human speciation via impacts on post-transcriptional modification of critical brain-related genes.

**Author summary:** Post-transcriptional modification has important functions in biological systems, but its role in evolution has long been a mystery due to the difficulty in predicting how a mutation could impact this process. Here we applied the newly developed deep learning model Seqweaver on human lineage mutations (HLM) to predict their effect on RNA-protein binding affinity profile (RBP). We found that only a very small number HLM are influential to RBP. Influential HLMs enriched in a set of conserved genes that are enriched in cells and functions critical for cognition and bipedal walking, and severe mutations on these genes cause the disruption of cognition (neurodevelopmental disorders) and bipedalism (ataxia). NTRK2 and ITPR1 had the most aggregated pieces of evidence of functional importance, pointing to an important role in cognition and bipedal walking. Taken together, our result demonstrated that there is a small number of human lineage mutations that modified the post-transcriptional modifications of conserved neuronal genes, which contributed to human speciation.

## Introduction

Human beings have long been interested in the question of what makes us unique from other animals[1]. Around 4% of the genome of Homo sapiens is different compared with our closest relative species chimpanzee (Pan troglodytes), including around 35 million single nucleotide variations and around 90 Mb regions with structural variations[1]. With the rapid development of sequencing and computational techniques, comparative genomics with other primates and archaic humans has also revealed novel genetic divergence[2]. These genetic differences cover almost all evolutionary events that distinguish humans from other primates, except the small number of extranuclear DNA[3]. Thus, researchers have conducted both biological[4] and computational[5] analyses to pinpoint the causal human lineage mutations that contributed to human speciation. These efforts provided valuable insights into evolution and biomedicine research, such as the discovery of the critical role of NOTCH2NL in neurogenesis and neurodevelopmental disorder[4].

However, the sparsity of influential human lineage mutations greatly challenged such analyses. Homo sapiens have undergone millions of years of purifying selection starting from our common ancestor, which would have eliminated the vast majority of mutations that could bring about deleterious consequences[6]. Thus, the remaining human lineage mutations would be mostly neutral. Based on this notion, the Combined Annotation-Dependent Depletion (CADD)[6] model directly used mutations that were fixed in human lineage as “proxy-neutral” in the training set, and used them to learn what features should a “neutral” mutation has. The high accuracy of CADD in predicting mutation deleteriousness supported their hypothesis that most human lineage mutations did not have any phenotypic consequence. Thus, the candidate gene analysis focusing on specific fixed mutations has a low prior probability of pinpointing the causal, influential human lineage mutations.

Another challenge lies in the functional analysis of non-protein-alternating mutations, which are nonsynonymous mutations whose mutation effects could not be directly estimated. Thus, most comparative genomic studies only focus on amino acid alteration, such as the ratio of nonsynonymous over synonymous coding mutations of a specific gene (dN/dS)[7]. Using dN/dS, a previous study[7] has revealed proteins that underwent significant positive selection during human speciation and their contribution to human cognition. However, theoretically speaking, human lineage mutation that did not alter amino acid could also contribute to phenotypic consequences via the alteration of transcription and post-transcriptional modification. Researchers have found evidence of the role of gene expression level[8] and alternative splicing[9] alterations in human evolution. Some technical innovations like RNA sequencing of human-chimp fusion cells[10] also provided a new opportunity to study non-coding mechanisms of human evolution. However, a systematic assessment of their role in human evolution is still lacking for post-transcriptional modification.

The newly developed deep learning model Seqweaver[11] provides a new opportunity to tackle these challenges. Seqweaver could take a DNA sequence as input and predict the binding affinity between the corresponding RNA sequence and 217 RNA-binding proteins. For a mutation, Seqweaver predicts RBP affinity for both reference and mutated sequence and takes their difference as the mutation’s impact on RBP binding. This prediction quantifies the human lineage mutations’ effect on post-transcriptional modification, enabling us to find the most influential mutations and systematically assess the role of post-transcriptional modification in human evolution. In this study (Figure 1), we applied Seqweaver to human lineage mutations and analyzed the signals of natural selection on them. Based on the existing knowledge stated above, we would like to answer the following questions in the current study: First, whether mutations with the largest impacts on RBP have been eliminated from human lineage. Second, whether of the small number of survived influential mutations have promoted human evolution by modifying post-transcriptional modification. Third, which highly influential mutations and target genes have important functions in Homo sapiens. We propose that such influential mutations could serve as an ideal candidate for future functional validation.

**Figure 1.**
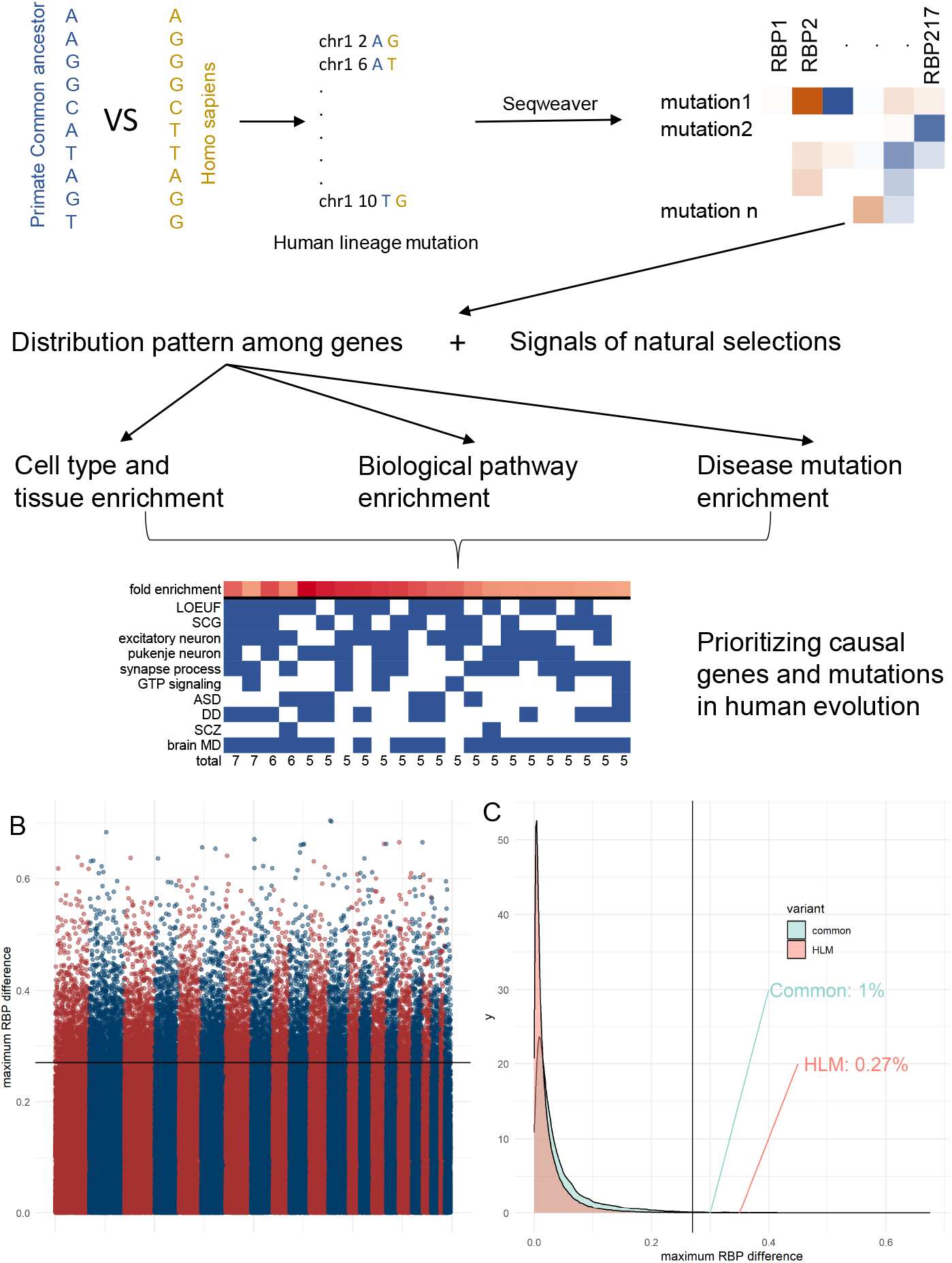
Overview of the study. A: study flowchart. B: Manhattan plot of Seqweaver result. Y axis denoted the maximum RNA-Binding profile difference (ΔRBP) of each human lineage mutation (HLM). X axis denoted the chromosome position of each HLM. C: distribution of ΔRBP.

## Result

### Strong negative selection on HLM’s impact on post-transcriptional regulation

We obtained a list of single nucleotide variations between primate common ancestor (multiple alignment) and human hg38 from CADD training set[10]. After the removal of variants within low-quality genomic regions and keeping only human-specific and fixed mutation (method), we retained 13,007,486 mutations for analysis, defined as human lineage mutation (HLM). As expected, these HLMs were strongly depleted in functionally important genomic regions, including exonic (Odds ratio, OR=0.49, p<10^− 100^), genic (including exon, intron and flanking regions, OR=0.90, p<10^−100^), and transcribed (OR=0.84, p<10^−100^) regions (Table S1). HLM were also depleted in tissue-specific active chromatin states (OR=0.76∼0.94 for 222 human tissues in the epimap database[12], Table S2) and cell type-specific open chromatin regions (OR=0.82∼0.96 for single-cell ATAC peak from 222 human cell types[13], Table S3). These results suggested that functional consequences of HLM were mostly intolerant and suffered strong negative selection.

Next, we set out to analysis HLM’s impact on post-transcription modification and extracted a list of 5,001,228 HLM falling within transcribed regions of coding genes,. By applying Seqweaver deep learning model[11], we quantified the impact of HLM on 217 RNA-binding protein affinity (ΔRBP, Figure 1B) and compared it with the common SNV of the human population in GnomAD[14] (Figure 1C). HLM generally had a significantly smaller ΔRBP than human common SNP (Wilcoxon test p<10^−100^). We calculated the maximum ΔRBP for each variant and found that only 13,475 transcribed HLM (0.27% of all HLM) had a maximum ΔRBP larger than the top 1% threshold of common SNP (fold change [FC]=0.27%/1%=0.27, Figure 1C), indicating a strong negative selection of HLM’s impact on RBP. Separately analyzing each of the 217 RBP profiles, we observed the strongest negative selection on ΔRBP related to alternative splicing (FC<0.01 for ΔRBP of TACA-spliced site binding, Table S4), indicating strict intolerance on RNA splicing alteration. To further validate this result, we directly quantified the impact of HLM and common SNP on alternative splicing by spliceAI[15] (Methods) and observed similar depletion of influential HLM, indicating purifying selection (FC=0.06, Figure S1).

Based on these results, we defined the 13,475 HLM with the largest ΔRBP as influential HLM and hypothesized that they might have an important role in human evolution. Top influential HLM falling on gene encoding histamine receptor H1 (max ΔRBP=0.75), as well as HLM on other brain-preferentially expressed genes like AMZ1 (max ΔRBP=0.78) and MPRIP (max ΔRBP=0.73). Interestingly, although HLM generally had small impact on RBP, these top influential HLM had larger impact than top common SNP: the largest max ΔRBP for common SNP was 0.70, smaller than these top HLM. We used these RBP-HLM for further functional research.

### Influential HLM enriched in genes highly conserved in primates and human

Next, we analyzed the biological significance of influential HLM. By applying a Poisson regression model (method), we found that influential HLM had a dramatically uneven distribution among protein-coding genes (Figure 2A). For example, some genes like USP25 carried six times more influential HLM than expected under null distribution, whereas 112 genes were expected to carry at least three influential HLM but actually carried none. Thus, we ranked all protein-coding genes according to the extent to which they enriched for influential HLM. We found that both top 10% (most enriched for influential HLM) and bottom 10% (most depleted for influential HLM) genes are more likely to be intolerant to loss-of function mutations[16] (Odds Ratio [OR]=2.16 and 1.70, Fisher test p=8.56×10^−26^ and 2.74×10^−12^, respectively, Figure 2B). They were also both more conserved in primate lineage[7] (Odds Ratio [OR]=1.53 and 1.42, Fisher test p=3.85×10^−15^ and 2.74×10^−10^, respectively, Figure 2C). One possible explanation of this result is that some sequences on some of the conserved genes are by nature more sensitive to mutation than other genes, and the mutations on these sequences are naturally more likely to be influential. To rule out this possibility, we applied saturated mutagenesis (Method) around HLM and found that random mutations on each gene did not have significant difference in the predicted effect on RBP profile (Figure S2).

**Figure 2.**
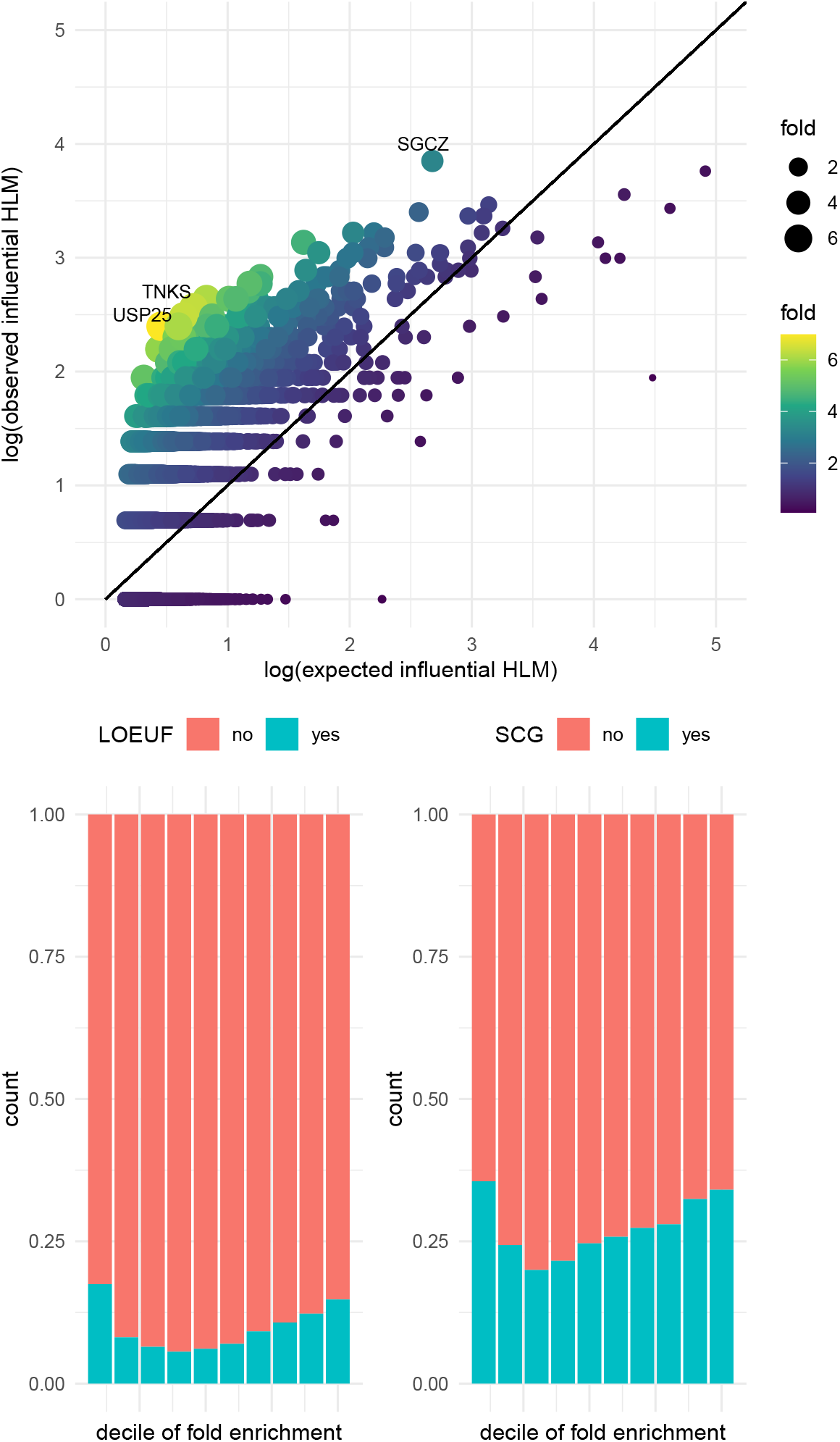
Gene-level analysis. A: observed vs expected number of influential human lineage mutations (HLM) on each gene. Color and dot size showed the fold enrichment of influential HLM. B: Each bar represented a decile of gene ranked by fold enrichment of influential HLM (the leftmost bar represented the most enriched decile). Bar color showed the proportion of loss-of-function intolerant genes within each decile. LOEUF: loss-of-function observed/expected upper bound fraction. C: same as B, but showing the proportion of primate lineage constraint genes defined by Dumas et al. SCG: selectively constraint genes.

This bimodal distribution of conservation is in contrast with the classic notion of gene conservation: mutations on essential and conserved genes are less likely to survive natural selection, thus we would observe a depletion of influential mutations on these genes. As a negative control, we ranked all genes according to the enrichment of influential common SNPs, and observed the expected unimodal distribution under this classic notion (Figure S3). Specifically, Genes more depleted for influential common SNP were more likely to be LoF-intolerant and conserved in primate lineage, in contrast to RBP-HLM enrichment results. We hypothesized that, instead of surviving natural selection, these influential RBP-HLM on essential genes were actually favored by natural selection and contributed to the evolutionary force that made us human.

To further verify this hypothesis, we defined 900 genes carrying at least one times more RBP-HLM than expected (fold enrichment >2), termed RBP-gene, for further functional analysis. As revealed above, RBP-gene significantly enriched in loss-of-function intolerant genes[16] (OR=2.10, Fisher test p=3.11×10^−14^), as well as primate constraint genes[7] (OR=1.42, Fisher test p=1.83×10^−6^). We reasoned that these genes were frequently hit by influential mutations during human speciation, thus might have an important role in human brain evolution and cognition function, and used them for downstream analysis.

### RBP-gene involved in synaptic functions and GTPase pathway

Human brain has undergone the most outstanding alteration during human evolution. Thus, if RBPgene truly contributed to human evolution, we would expect to see that these genes played critical roles in brain functions and should be highly expressed in neuron. Indeed, by analyzing Single-cell transcriptome data[17], we found that RBP-gene were highly expressed in central nervous system (CNS): in fetal tissue, RBP-gene were only enriched in cerebellum, including Pukinje cells and several other subtypes of neurons (p<10^−100^, Figure 3A). RBP-gene that highly expressed in Pukinje neuron included USP25, ITPR1, KCNH8 and SCN8A, etc. In adult tissue, RBP-gene enriched in different subtypes of excitatory neuron from visual cortex and frontal cortex (p<10^− 100^, Figure 3B). RBP-gene that highly expressed in cortex excitatory neuron included NTRK2, NLGN1, GABRB2 and CACNA1D, etc. The number of CNS cell types with nominally significant enrichment (29) were also larger than all other systems and organs. Interestingly, cerebral cortex and cerebellar Pukinje cells are vital for cognition functions and cooperation of bipedal walking[7], both of which are the key functions during human evolution[7]. Taken together, RBP-gene were mostly enriched in fetal cerebellum and adult cortex, since both the enrichment p value and the number of enriched cell types were the highest compared to other tissues and organs.

**Figure 3.**
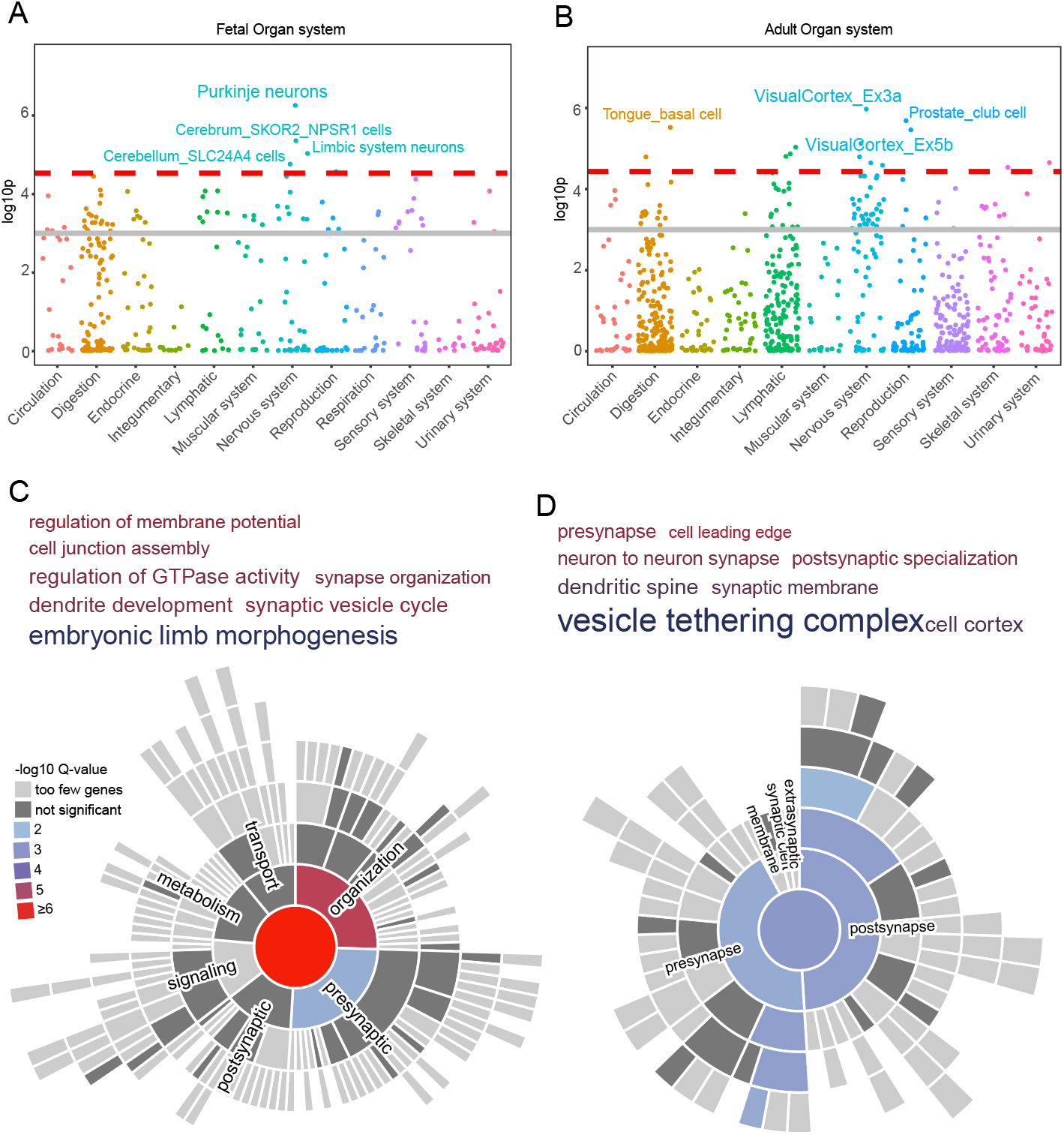
Functional characteristics of RBP-gene. A & B: WebCSEA results of cell type-specific expression in different embryonic (A) and adult (B) tissues. Low p-value indicated that RBP-gene specifically expressed in the cell type at high significance. C & D: Gene ontology enrichment analysis of biological process (C) and cellular component (D). Font size indicated fold enrichment, and color indicated enrichment p-value. We manually selected specific terms with FDR-adjusted p<0.05. E & F: synGO enrichment analysis of biological process (E) and cellular component (F). Color indicated enrichment p-value.

We further analyzed the biological functions of RBP-gene by gene ontology (GO) analysis. As shown in Figure 3C and Table S5, RBP-gene significantly enriched in synapse organization (p=6.94×10^−7^), regulation of membrane potential (p=1.37×10^−6^), dendrite development (p=1.33 × 10^−6^), synaptic vesicle cycle (p=1.32 × 10^−5^), cell junction assembly (p=1.88×10^−6^) as well as other pathways related to neuronal and synaptic functions. Despite neuron-related pathways, RBP-gene also showed strong enrichment in regulation of GTPase activity (p=5.57×10^−10^). In cellular component analysis (Figure 3D), we found that RBP-gene mainly located in various component of neuron, including pre-synapse (p=1.31×10^−20^), postsynaptic specialization (p=3.63×10^−20^) and dendritic spine (p=1.49×10^−13^).

Since all these results indicated the important role of RBP-gene in synaptic functions, we further applied synGO[18] pathway analysis, an expert-curated synaptic ontology database, to refine the result. Consistently, synGO analysis also confirmed that RBP-gene were enriched in synapse process (p=9.56 × 10^−9^) and synapse organization (p=2.54×10^−7^), as well as their child terms (Figure 3E). For localization analysis, synGO revealed that RBP-gene involved in both pre- and post-synaptic complex (adjusted p=2.75×10^−8^ and 6.62×10^−10^, respectively) as well as moderate enrichment in the sub-regions (pre-synaptic activate zone and post-synaptic density; Figure 3F). Taken together, genes involved in synaptic organization and other pathways of synapse carried excess number of HLM that had large impact on post-transcriptional modification, which might contribute to the evolution of human brain.

### RBP-gene carried excess severe mutations of neurodevelopmental disorders

Human brain evolution has shaped the cognitive functions of modern human, and contributed to the genetic basis of brain disorders. we hypothesized that rare, damaging variants that disrupt RBP-gene have contribution to brain disorders. As shown in Figure 4A, using published cross-sectional burden test[19] result of Autism, we found that compared with background genes, RBP-gene generally carried excess burden of damaging coding mutations in patients (fold enrichment=4.33, Fisher test p=1.19×10^− 5^). Similar but less significant result was also obtained for schizophrenia[20] (fold enrichment=4.58, Fisher test p=0.004). In trio-based WES analysis, RBPgene also carried excess de novo damaging mutations in autism probands[21] (fold enrichment=2.87, Fisher test p=0.0002). and developmental delay probands[22] (fold enrichment=2.01, Fisher test p=9.01×10^−6^). Similarly, RBP-gene were also more likely to be the risk genes of Mendelian brain disorder (fold enrichment=1.34, Fisher test p=0.001). We repeated these analyses after controlling covariates like LOEUF and gene length (Method), and got consistent significant results albeit with lower statistical power (Table S6). These results suggested that rare and severe mutations on RBP-gene are more likely to cause neurodevelopmental disorders.

**Figure 4.**
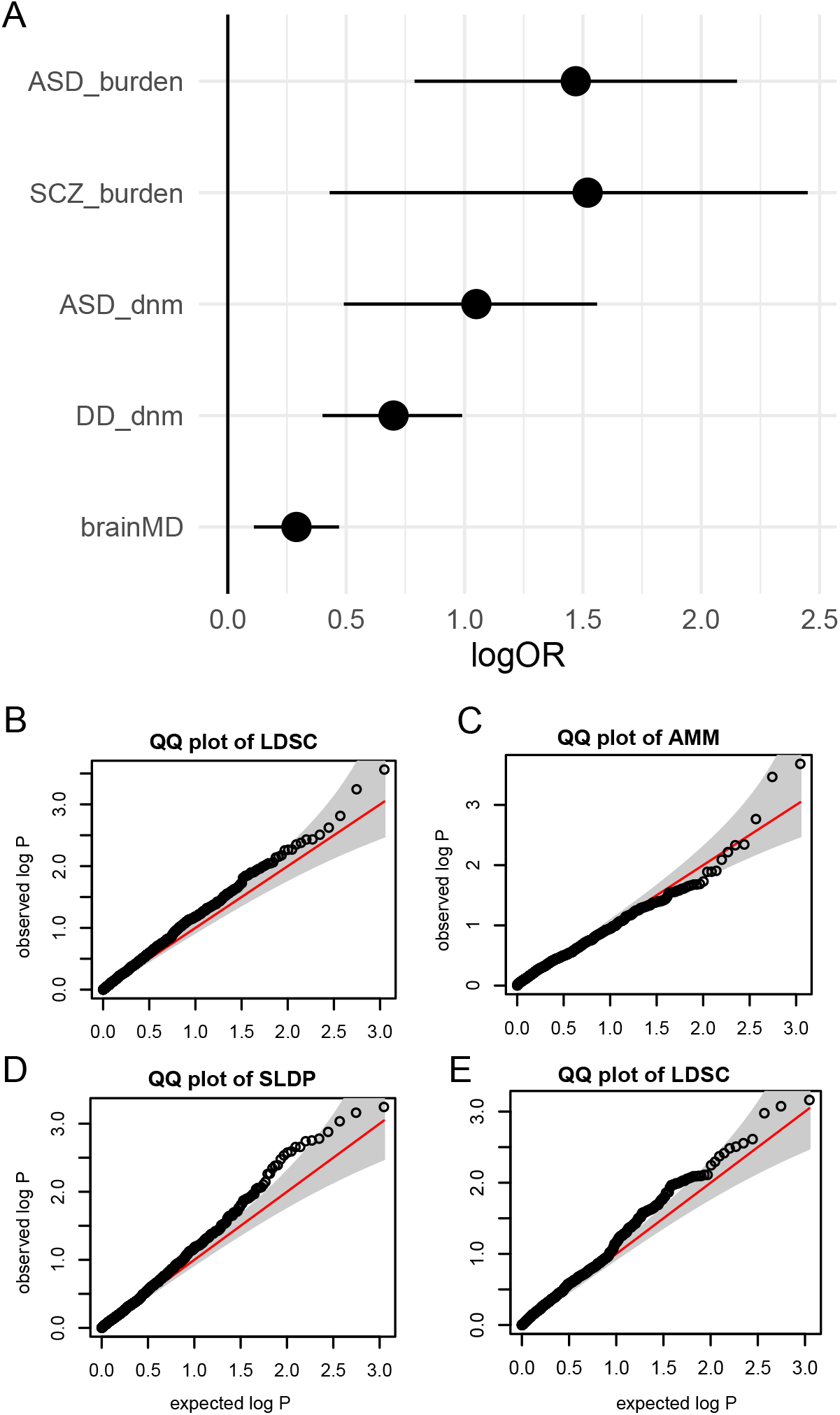
Phenotypic consequence of RBP-gene. A: Enrichment of RBP-gene in brain diseases gene lists. Error bar indicated 95% confidence interval. Dnm: de novo mutation. brainMD: brain Mendelian disorders. B: quantile-quantile plot of LDSC p-value for trait heritability enrichment around RBP-gene. C: quantile-quantile plot of Abstract Mediation Model (AMM) p-value for trait heritability enrichment around RBP-gene. D: quantile-quantile plot of sign linkage disequilibrium profile (SLDP) p-value for trait heritability enrichment around humanization score. E: quantile-quantile plot of LDSC p-value for trait heritability enrichment around absolute humanization score.

We further analyzed whether common variants on RBP-genes had a significant phenotypic consequence. By applying linkage disequilibrium score regression (LDSC) on a set of about 1,000 polygenic traits, we found that SNPs around RBP-gene did not explain significantly higher proportion of trait heritability (FDR-adjusted p value > 0.05, Figure 4B and Table S6). Using abstract mediation model (AMM, method) instead of LDSC also revealed no significant result (Figure 4C and Table S7). This result could be expected if the effect of post-transcriptional modification on human evolution is oligogenic instead of polygenic. In fact, if human speciation is driven by a few vital mutations on a few vital genes, there would not be a large number of variations with a small but non-zero contributions. Under this scenario, the large number of common SNP actually had no association with post-transcriptional modification during human speciation. Given the fact that influential mutations are mostly eliminated by purifying selection and the remaining RBP-HLM is very sparse, the oligogenic view is plausible.

### Oligogenic view of post-transcriptional modification changes in human evolution

To assess this theory, we used Seqweaver to calculate RBP difference of all genome-wide transcribed regions between hg38 and primate common ancestor genome alignment (Method). We then applied Seqweaver to calculate how each 1000 genome[23] common SNP intensify (over-humanize) or weaken (de-humanize) this difference, termed humanization score. If human evolution on RBP profile is polygenic, a large number of transcribed regions would have RBP alterations that have small phenotypic effect. Then, common SNPs with large over- or de-humanization effect would collectively explain excess proportion of trait heritability. However, this is not true: humanization score was not significantly associate with trait heritability in LDSC (all trait had FDR-adjusted p value >0.05, Figure 4E and Table S8); the result is still insignificant when taking direction of humanization score into consideration by using sign linkage disequilibrium profile (SLDP; Method) (Figure 4D and Table S9). Taken together, these findings were in line with an oligogenic view of human evolution of RBP profile, although it is difficult to draw statistical conclusion from null result.

### Prioritizing ITPR1 and NTRK2 in human evolution

The oligogenic view of human evolution of RBP profile suggested that among the 900 RBP-genes showing enrichment of influential HLM, only a small subset might actually contribute to human evolution, which gave rise to the functional enrichment of RBP-gene. Thus, we sought to aggregate all functional and phenotypic evidences in the above analysis for all genes (Table S10), and prioritize the most probable genes that took part in the human evolution of RBP profile. As shown in Figure 5A, there were 22 RBP-gene that had at least five evidences of functional and phenotypic importance. Two top genes, NTRK2 and ITPR1, had seven aggregated evidences.

**Figure 5.**
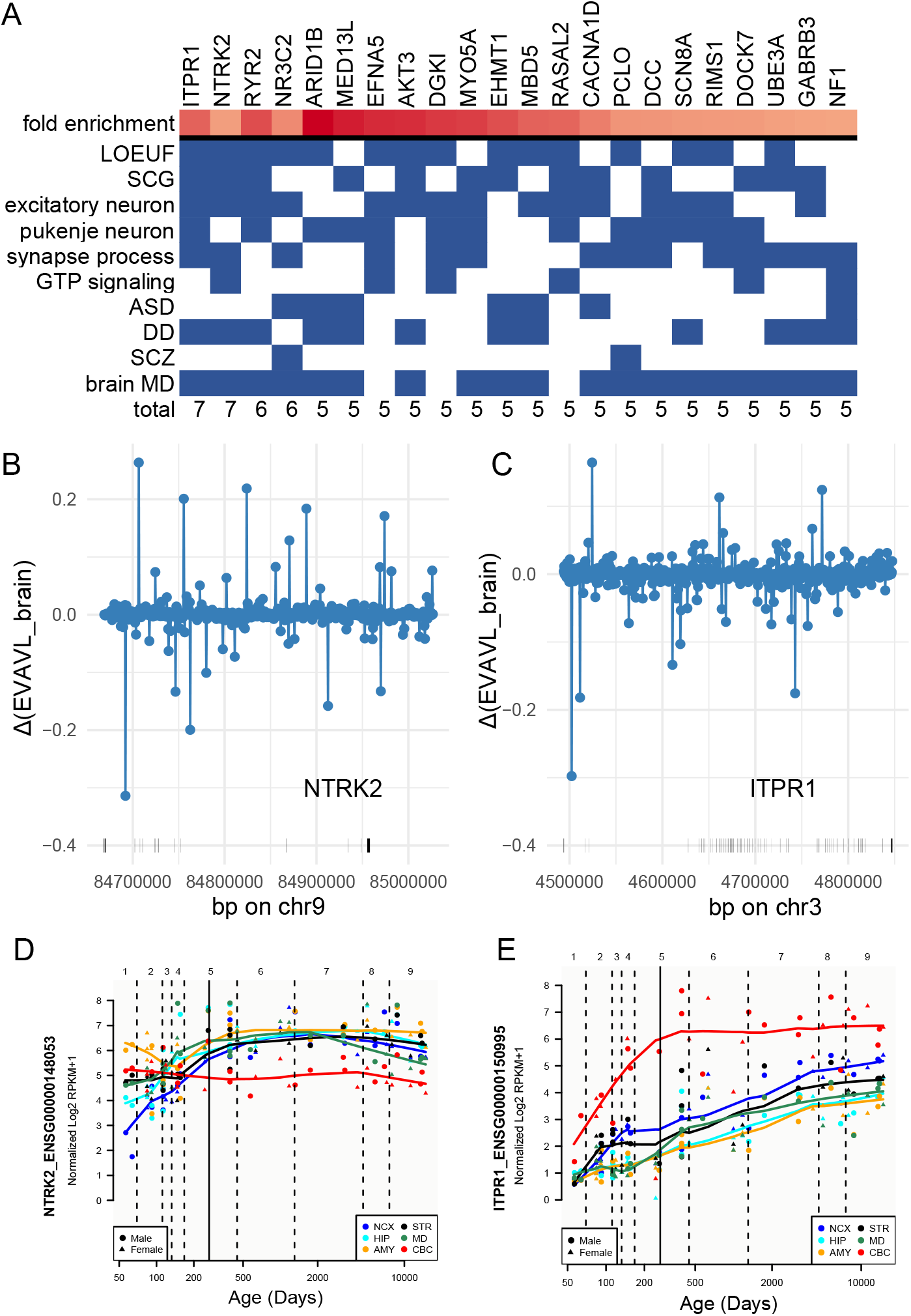
Prioritizing top candidate genes. A: All RBP-genes with at least five aggregated evidences. B: Seqweaver predicted difference of EVEVL-RNA binding affinity in human brain between hg38 and primate common ancestor sequence of NTRK2. Each dot represented a 1000bp block on NTRK2. C: same as B, but for ITPR1. D: spatiotemporal expression of NTRK2 in human developing brain. Figure generated at Brainspan website. E: same as D, but for ITPR1.

NTRK2 (chr9: 84668522-85027054) encodes neurotrophic receptor tyrosine kinase 2. NTRK2 is intolerant to LoF in human and is conserved in primate lineage, highly expressed in excitatory neuron and takes part in synapse process and GtP pathway (Figure 5A). Seqweaver revealed that HLM at the forth intron of NTRK2 has profoundly decreased the binding affinity with EVAVL in the human brain (ΔRBP=- 0.33, Figure 5B), which was the largest alteration among all 217 RBP. In Brainspan data[24], cerebral expression level of NTRK2 consistently increased until childhood, and remained at the peak expression level until adult (Figure 5D). This is in line with the fact that NTRK2 is associated with neurodevelopmental disorders including developmental delay and multiple Mendelian brain disorders (Figure 5A) like Astrocytoma and Developmental and Epileptic Encephalopathy[25].

ITPR1 (chr3: 4493348-4847506) encodes inositol 1,4,5-trisphosphate receptor type 1. ITPR1 is also conserved in both human and primate lineage, and has high expression in both excitatory neurons and Purkinje neurons (Figure 5A). In Seqweaver analysis, HLM in the second intron of ITPR1 caused the most profound alteration of RBP (ΔRBP=-0.30, Figure 5C). Interestingly, this alteration was also on EVAVL binding affinity in the human brain, just like NTRK2. Consistent with its high expression in Purkinje neurons, ITPR1 has the highest expression in the cerebellum, and the expression value continuously increases throughout human developmental periods (Figure 5E). Furthermore, ITPR1 has been identified as a risk gene for several cerebellar genetic disorders, such as different subtypes of Spinocerebellar Ataxia and Gillespie Syndrome[25], suggesting that ITPR1 may have played a role in the evolution of bipedal walking.

## Discussion

In this study, we applied a deep learning model Seqweaver on genome-wide human lineage mutations to predict their impact on post-transcriptional modification. We found that such impact is highly intolerant in the human lineage, and a small number of influential HLM has enriched on a set of conserved genes that had both functional and phenotypic significance. We inferred that the cis-regulation of post-transcriptional modification on this set of conserved genes has contributed to human evolution.

The major evidence of this conclusion is the bimodal relationship between influential HLM enrichment and gene conservation, as shown in Figure 2. Using dN/dS metric of coding mutations, a previous study[7] has demonstrated that genes carrying an excess number of influential coding mutations during human speciation have undergone a positive selection and are key genes of human evolution. Expanding this view to non-coding regions, the study of human accelerating regions[26] found that sequences that are conserved across species but carry excess HLM are vital for human brain expansion. In line with these studies, we also found that while conserved genes were generally depleted for HLM influential on post-transcriptional modification, there were a subset of conserved genes carrying unexpectedly large number of influential HLM. Our results together with previous findings revealed that influential HLM on conserved genes and regions contributed to the positive selection that made us human, via alterations of both protein sequence, transcription regulation and post-transcriptional modification.

Furthermore, the functional and phenotypic characteristics of this set of RBP-genes also supported their role in human evolution. Cortical expansion and cerebellar reorganization are the critical steps for evolving cognition[27] and bipedal walking[28], two major characteristics of modern human. In our analysis, RBP-genes were highly expressed in excitatory neuron and Pukinje neuron of cortex and cerebellum. The rare and severe mutation on them were also associated with both polygenic and Mendelian neurodevelopmental disorder characterized by deficiency in cognition and bipedalism (ataxia). Given the fact that RBP-genes are conserved, carried excess number of influential HLM, and are associated with cognition and bipedalism, it is plausible to state that they had important roles in human evolution.

Among these RBP-genes, we prioritized NTRK2 and ITPR1 as the potential key genes of human evolution. Our result showed that they both carried excess HLM that had large impact on post-transcriptional modification, and are both conserved in human and primate lineage. NTRK2 encodes neurotrophic tyrosine kinase receptor type 2, a receptor that can be activated by multiple neurotrophins and regulated downstream neuronal proliferation, differentiation, and neurotransmitter systems[29]. NTRK2 could also regulated astrocyte proliferation via Rho-GTPase system[30], and is associated with multiple neuropsychiatric disorders[29,31]. It could be inferred that NTRK2 is a key gene in cortex development and human cognition. ITPR1, on the other hand, mainly express in Pukinje cells and control the calcium release by binding Inositol 1,4,5-triphosphate. Deficiency of ITPR1 in mice model led to motor discoordination[32], and ITPR1 mutations in human were associated with various types of ataxia[33]. Thus, TIPR1 could have an important role in coordination of bipedal walking during evolution.

It should be noted that since Seqweaver was trained on human RBP data but not on other hominin’s RBP data, our study relied on the assumption that positive selection on post-transcriptional modification only happened in cis-manner instead of trans-manner. That is, only the HLMs on RNA have made an effect, but the entire RNA binding protein system itself was constant across species. We found two pieces of evidence to support this assumption. Firstly, there are Seqweaver models that were trained on mouse RBP data, and these models are proved valuable in predicting pathogenic mutations of autism[11,34], which suggested that cross-species differences in RBP system did not drive a systematic bias. Secondly, our result showed that (Table S10) RNA binding proteins like RBFOX1 were among the genes that were most depleted from influential HLM, further supported that RBP themselves were highly conserved and were absent from significant alterations during evolution.

Our study has other limitations. Structural variations have been shown to play an important role in human evolution[35], but current sequence-based deep learning models like Seqweaver are unable to evaluate their effects. The lack of statistical tests on Seqweaver estimation has also obstructed us from stating any particular HLM to be confidentially influential. Instead, we could only analyze the overall patterns of influential HLM, and prioritize some top HLM and genes as the most probable causal mutations and genes. Future experimental validations on the top HLM and RBP-genes will help to fill in this gap.

In conclusion, we demonstrated that despite the strong purifying selection on human lineage mutations, there is a small number of HLM that had a large impact on post-transcriptional modification of essential genes and contributed to human evolution. These essential genes take part in synaptic functions and neurodevelopmental disorders and are ideal candidates for future analysis.

## Method

### Data preprocessing and characterization

We downloaded the list of human lineage mutations from CADD training set[6], which was obtained by comparing hg38 against Enredo, Pecan, and Ortheus (EPO) 6 primate alignment[36] of the common ancestor. We removed HM within low-quality regions of hg38 (gap region defined in UCSC genome browser[37]), including short arm gaps, heterochromatin gaps, telomere gaps, gaps between contigs in scaffolds, and gaps between scaffolds in chromosome assemblies. We also excluded HLM within centromere regions[38]. We then obtained a list of single nucleotide differences between humans and chimpanzees (hg38 vs. Pantro5) from Gokhman et al.[10] and took the intersection to remove mutations that were not specific to the human lineage.

For Seqweaver analysis, we retained only HLM that fell on the transcription regions of coding genes, where the transcription start and end site were obtained from Ensembl database, as provided by the Seqweaver toolkit.

We analyzed whether the overall HLMs were enriched in or depleted from the following genomic regions:

1. exonic, genic, and transcribed regions, downloaded from UCSC genome browser.
2. active chromatin regions of 222 tissues and cell types: we downloaded from epimap database[12] the chromHMM[39] 18-chromatin state annotations of each sample. We defined the following annotations as “active chromatin regions”: “TssA”, “TssFlnk”, “TssFlnkU”, “TssFlnkD”, “Tx”, “EnhG1”, “EnhG2”, “EnhA1”, “EnhA2”. We grouped all samples according to tissue names and embryo/adult status, leading to 222 groups in total. Within each group, we kept all genomic regions that were marked as “active chromatin regions” in at least half of the samples.
3. Open chromatin regions of 222 cell types: We downloaded the single-cell ATAC-seq peak annotation from Zhang et al.[13], which consisted of 222 cell types covering both prenatal and postnatal cells from all parts of the body.

For all of these annotations, we excluded gap regions before analysis. HLM were mapped to each annotation by bedtools[40]. For each annotation, we summed up the total length and calculated the expected number of HLM on them. We then applied a binomial test to see if observed number of HLM was significantly differed from expected number.

### Seqweaver analysis

Seqweaver was applied at default settings[11]. Only 217 models on human RBP were applied, and mouse RBP model were excluded. To define the threshold of influential HLM, we obtained a list of common SNP in human population, which had minor allele count > 10,000 in GnomAD[41] v3.1.2 whole-genome sequencing data (non-neural disorder subset), and applied Seqweaver on them. We also excluded SNP within gap regions as defined above. For each of the 217 RBP, we calculated the top 1% threshold of absolute value of predicted RBP binding affinity (ΔRBP) difference of all common SNP. We also calculated the top 1% threshold of maximum absolute ΔRBP. HLM that had maximum ΔRBP larger than this threshold were considered influential HLM.

We applied a saturated mutagenesis analysis by generating all possible SNP within 200 bp windows around each HLM and input them to Seqweaver. We assigned each of these generated SNP to genes and tested if the overall maximum ΔRBP of generated SNP in each gene group significantly differed from each other.

For sequence level Seqweaver analysis, we first downloaded the EPO primate common ancestor alignment (corresponded to hg38) from Ensembl[36]. For each gene, we used a sliding window of 1000 bp and 500 bp per step size to cover its full length, and extracted DNA sequence of hg38 and common ancestor alignment for each of the 1000 bp-length blocks. These sequences in fasta format were input to Seqweaver at sequence mode. We calculated the difference of 217 RBP binding affinity between hg38 sequence and ancestor sequence for each block.

### spliceAI analysis

spliceAI[15] is a deep learning tool that predict the probability that a mutation could influence acceptor gain, acceptor loss, donor gain and donor loss of the closet splice site. We downloaded the masked prediction result of spliceAI from Illumina website, extracted results for both HLM and common SNP, and calculated the maximum score of each variant. We also calculated the top 1% threshold of common SNP similar to Seqweaver analysis.

### Gene level analysis

Taking all 17,329 protein coding genes together, we applied the following Poisson regression with log link to estimate the expected number of influential HLM on each gene:

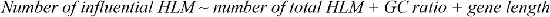

To avoid log (0), we added one pseudo count to all genes. We took the predicted value from this regression as the expected number of influential HLM, and used the ratio of observed to expected number as the fold enrichment. We ranked and grouped all gene into deciles in the descending order of fold enrichment, and calculated the proportion of two sets of genes in each decile: firstly, human conserved genes were defined as top 10% of genes in GnomAD LOEUF score[14]; secondly, primate conserved genes were defined as genes with the lowest 25% dN/dS across six primates calculated by Dumas et al.[7]. The significance of enrichment of these gene lists in each decile was calculated by Fisher test. We also repeated these analyses on common SNP as a negative control.

### Functional analysis

We defined genes with fold enrichment > 2 as RBP-genes and used them for functional analysis. We used WebCSEA tool to apply expression enrichment test on a comprehensive set of published single-cell RNA sequencing data covering different embryo and adult tissues. The permutation-based combined p-value for each cell types were used to define significantly enriched cell types. For the highlighted cell types, we extracted top 5% genes showing specific expression, as defined by t-statistics calculated by WebCSEA. For highlighted genes, we additionally extracted their expression trajectory among brain development using online tool of Brainspan[24].

We used ClusterProfiler[42] R package to conduct Gene Ontology (GO) Biological Process (BP) and Cell Component (CC) enrichment analysis. We only retained pathways with >10 and <500 genes for analysis. Background genes were defined as all genes with GO annotation. We applied simplify() function to remove similar pathways (highly overlapped or child-parent term of every other). We reported pathways with FDR-corrected p value<0.05. We also applied SynGO[18] enrichment analysis, with similar settings except that the background gene list was defined as all brain-expressed genes.

To analyze whether RBP-genes were significantly associated with neuro-developmental disorders, we collected disease genes from the following resource:

1. cross sectional autism whole-exome sequencing (WES) data (11,986 cases, 23,598 control)[19]. All genes with FDR-adjusted p-value of transmitted and de novo association (TADA)[43] < 0.05 were collected.
2. Combined Schizophrenia cross sectional WES data (24,248 cases, 97,328 controls) and trio WES data (3,402 trios)[20]. Significance threshold was FDR-adjusted p-value of meta-analysis < 0.05.
3. trio-based WES data of developmental delay (31,058 trios). We collected genes with FDR-adjusted p-value of DeNovoWEST <0.05.
4. family-based WES data (15,306 probands) of autism[21]. We collected genes with FDR-adjusted p-value of DeNovoWEST <0.05.
5. risk genes of brain Mendelian disorders. We downloaded the gene-disease-organ association tables from Gene ORGANizer[44] database, and retained only genes associated with brain with high confident.

We tested whether RBP-gene enriched in these gene sets by Fisher test. To control potential bias, for each gene set we additionally ran a Logistic regression on all included genes:

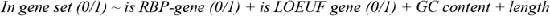

to verify the enrichment result. Positive regression coefficient of term *is RBP-gene (0/1)* was considered evidence of enrichment, and its p-value was used to evaluate the significance.

### Heritability enrichment analysis of polygenic traits

As previously described[45], we uniformly collected and pre-processed a set of GWAS summary statistics that 1) came from European ancestry; 2) SNP heritability h^2^>0.01; 3) z score of h^2^ >4; 4) sample size >10,000. We applied Linkage Disequilibrium Score Regression (LDSC)[46] to analyze whether heritability of these traits enriched in common SNP around RBP-gene (window size = 100kb). We used 1000 Genome[23] European population as reference panel, only SNP within hapmap3[47] project were included, and the baseline model and other parameters of LDSC were set at default. To control the bias of incorrect SNP-to-gene mapping, we additionally applied Abstract Mediation Model (AMM)[48], an extension of LDSC that also considered the k-nearest genes of each SNP. We used the default hyperparameters provided by AMM, which were estimated by the benchmark gene set of loss-of-function intolerant genes. We directly transformed the enrichment z score of AMM into p-value under normal distribution and used for FDR adjustment, without log-transformation of the enrichment.

To analyze whether the human-specific directional impact of common SNP on RBP profile has a phenotypic consequence, we first calculated a “humanization score” of each 1000 Genome common SNP. As described above, we first used 13,520,465 overlapping blocks (1,000 bp length each) *b*=1,2,…13,520,465 to cover full length of all protein-coding genes. For each block *b*, we used sequence-mode Seqweaver to calculate a vector of RBP for hg38 sequence on *b* (*hg*38_*b*_(*r*), *r* = 1,2, … 217), a vector of RBP for common primate ancestor sequence on *b* (*anc*_*b*_(*r*)), and calculated the difference of them *ΔRBP*_*b*_(*r*) = *anc*_*b*_(*r*) − *hg*38_*b*_(*r*). We then mapped each 1000G common variation *s* to a block *b* = *f*(*s*), whose midpoint was closest to *s* (note that multiple SNP could be mapped to a same block). If *s* was not on a gene body, *f*(*s*) = 0. We generated a DNA sequence corresponded to *s* by SAMtools[49] consensus option, where the reference fasta was the sequence of block *b* = *f*(*s*). We applied Seqweaver on this DNA sequence to obtained vector *RBP*_*f*(*s*)_(*r*), and calculated the humanization score (HS) of SNP *s* as the inner product of two vectors:

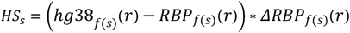

Negative value of *HS*_*s*_ indicated that SNP *s* modified human RBP profile in the opposite direction that all HLM on block *f*(*s*) collectively modified ancestor RBP profile, and it made human profile closer to ancestor profile, termed “de-humanization”. Likewise, Positive *HS*_*s*_ indicated that *s* modified human RBP profile further from ancestor RBP profile, corresponded to “over-humanization”. Our assumption is that: if human evolution on RBP profile is polygenic, then there will be a large number of block *b* whose *ΔRBP*_*b*_(*r*) had a small but non-zero phenotypic association. Then, SNP on these blocks that impact *ΔRBP* would also slightly impact phenotypes. Thus, *HS*_*s*_ would be associated with SNP-based trait heritability enrichment. We used two approaches to evaluate this association:

1. signed linkage disequilibrium profile regression (SLDP)[50], an extension of LDSC that accounts for the direction of each genomic annotations. Significant positive SLDP regression coefficient on HS indicated that over-humanization SNP collectively explain excess heritability of trait, and vice versa. Data preprocessing was the same as LDSC, and all parameters were set at default.
2. LDSC analysis on absolute HS value: we directly included absolute HS as a continuous annotation in LDSC. Significant positive LDSC regression coefficient indicated that SNP with relatively large RBP impact in regions with human-specific RBP profiles collectively explain excess heritability of trait.

## Acknowledgement

This work was supported by grants from the 2030 Science and Technology Innovation Key Program of Ministry of Science and Technology of China (No. 2022ZD020910001), the National Natural Science Foundation of China (No. 81971292, 82150610506) and the Natural Science Foundation of Shanghai (No. 21ZR1428600), the Medical-Engineering Cross Foundation of Shanghai Jiao Tong University (No. YG2022ZD026).

## Author Contribution

W.S and G.N.L designed the study. W.S collected the data, performed analysis and drafted the paper. S.Y and M.Z interpreted the result. All authors read, revised and approved the manuscript.

## Reference

1. Varki A, Altheide TK. Comparing the human and chimpanzee genomes: searching for needles in a haystack. Genome Res. 2005;15: 1746–1758. doi:10.1101/GR.3737405

2. Yousaf A, Liu J, Ye S, Chen H. Current Progress in Evolutionary Comparative Genomics of Great Apes. Front Genet. 2021;12: 1436. doi:10.3389/FGENE.2021.657468/BIBTEX

3. Lan YY, Heather JM, Eisenhaure T, Garris CS, Lieb D, Raychowdhury R, et al. Extranuclear DNA accumulates in aged cells and contributes to senescence and inflammation. Aging Cell. 2019;18. doi:10.1111/ACEL.12901

4. Fiddes IT, Lodewijk GA, Mooring M, Bosworth CM, Ewing AD, Mantalas GL, et al. Human-Specific NOTCH2NL Genes Affect Notch Signaling and Cortical Neurogenesis. Cell. 2018;173: 1356–1369.e22. doi:10.1016/j.cell.2018.03.051

5. Liu J, Robinson-Rechavi M. Robust inference of positive selection on regulatory sequences in the human brain. Sci Adv. 2020;6: 9863–9890. doi:10.1126/SCIADV.ABC9863

6. Kircher M, Witten DM, Jain P, O’Roak BJ, Cooper GM, Shendure J. A general framework for estimating the relative pathogenicity of human genetic variants. Nat Genet. 2014;46: 310–315. doi:10.1038/ng.2892

7. Dumas G, Malesys S, Bourgeron T. Systematic detection of brain proteincoding genes under positive selection during primate evolution and their roles in cognition. Genome Res. 2021;31: 484. doi:10.1101/GR.262113.120/-/DC1

8. Fraser HB. Gene expression drives local adaptation in humans. Genome Res. 2013;23: 1089–1096. doi:10.1101/gr.152710.112

9. Xiong J, Jiang X, Ditsiou A, Gao Y, Sun J, Lowenstein ED, et al. Predominant patterns of splicing evolution on human, chimpanzee and macaque evolutionary lineages. Hum Mol Genet. 2018;27: 1474–1485. doi:10.1093/HMG/DDY058

10. Gokhman D, Agoglia RM, Kinnebrew M, Gordon W, Sun D, Bajpai VK, et al. Human-chimpanzee fused cells reveal cis-regulatory divergence underlying skeletal evolution. Nat Genet. 2021;53: 467. doi:10.1038/S41588-021-00804-3

11. Park CY, Zhou J, Wong AK, Chen KM, Theesfeld CL, Darnell RB, et al. Genome-wide landscape of RNA-binding protein target site dysregulation reveals a major impact on psychiatric disorder risk. Nat Genet. 2021;53: 166. doi:10.1038/S41588-020-00761-3

12. Boix CA, James BT, Park YP, Meuleman W, Kellis M. Regulatory genomic circuitry of human disease loci by integrative epigenomics. Nature. 2021;590: 300–307. doi:10.1038/s41586-020-03145-z

13. Zhang K, Hocker JD, Miller M, Wang A, Preissl S, Correspondence BR, et al. A single-cell atlas of chromatin accessibility in the human genome. Cell. 2021;184: 1–17. doi:10.1016/j.cell.2021.10.024

14. Karczewski KJ, Francioli LC, Tiao G, Cummings BB, Alföldi J, Wang Q, et al. The mutational constraint spectrum quantified from variation in 141,456 humans. Nature. 2020;581: 434–443. doi:10.1038/s41586-020-2308-7

15. Jaganathan K, Kyriazopoulou Panagiotopoulou S, McRae JF, Darbandi SF, Knowles D, Li YI, et al. Predicting Splicing from Primary Sequence with Deep Learning. Cell. 2019;176: 535–548.e24. doi:10.1016/J.CELL.2018.12.015

16. Karczewski KJ, Francioli LC, Tiao G, Cummings BB, Alföldi J, Wang Q, et al. Variation across 141,456 human exomes and genomes reveals the spectrum of loss-of-function intolerance across human protein-coding genes. bioRxiv. 2019; 531210. doi:10.1101/531210

17. Dai Y, Hu R, Liu A, Cho KS, Manuel AM, Li X, et al. WebCSEA: web-based cell-type-specific enrichment analysis of genes. Nucleic Acids Res. 2022;50: W782–W790. doi:10.1093/NAR/GKAC392

18. Koopmans F, van Nierop P, Andres-Alonso M, Byrnes A, Cijsouw T, Coba MP, et al. SynGO: An Evidence-Based, Expert-Curated Knowledge Base for the Synapse. Neuron. 2019;103: 217–234.e4. doi:10.1016/j.neuron.2019.05.002

19. Satterstrom FK, Kosmicki JA, Wang J, Breen MS, De Rubeis S, An JY, et al. Large-Scale Exome Sequencing Study Implicates Both Developmental and Functional Changes in the Neurobiology of Autism. Cell. 2020;180: 568–584.e23. doi:10.1016/j.cell.2019.12.036

20. Singh T, Poterba T, Curtis D, Akil H, Al Eissa M, Barchas JD, et al. Rare coding variants in ten genes confer substantial risk for schizophrenia. Nature. 2022;604: 509–516. doi:10.1038/s41586-022-04556-w

21. Fu JM, Satterstrom FK, Peng M, Brand H, Collins RL, Dong S, et al. Rare coding variation provides insight into the genetic architecture and phenotypic context of autism. Nat Genet. 2022;54: 1320–1331. doi:10.1038/s41588-022-01104-0

22. Kaplanis J, Samocha KE, Wiel L, Zhang Z, Arvai KJ, Eberhardt RY, et al. Evidence for 28 genetic disorders discovered by combining healthcare and research data. Nature. 2020;586: 757–762. doi:10.1038/s41586-020-2832-5

23. 1000 Genomes Project Consortium. A global reference for human genetic variation. Nature. Nature Publishing Group; 2015. pp. 68–74. doi:10.1038/nature15393

24. Li M, Santpere G, Imamura Kawasawa Y, Evgrafov O V., Gulden FO, Pochareddy S, et al. Integrative functional genomic analysis of human brain development and neuropsychiatric risks. Science (80-). 2018;362: eaat7615. doi:10.1126/science.aat7615

25. Hamosh A, Scott AF, Amberger JS, Bocchini CA, McKusick VA. Online Mendelian Inheritance in Man (OMIM), a knowledgebase of human genes and genetic disorders. Nucleic Acids Res. 2005;33. doi:10.1093/NAR/GKI033

26. Doan RN, Bae B Il, Cubelos B, Chang C, Hossain AA, Al-Saad S, et al. Mutations in Human Accelerated Regions Disrupt Cognition and Social Behavior. Cell. 2016;167: 341–354.e12. doi:10.1016/j.cell.2016.08.071

27. Wei Y, de Lange SC, Scholtens LH, Watanabe K, Ardesch DJ, Jansen PR, et al. Genetic mapping and evolutionary analysis of human-expanded cognitive networks. Nat Commun. 2019;10: 1–11. doi:10.1038/s41467-019-12764-8

28. Jahn K, Deutschländer A, Stephan T, Kalla R, Wiesmann M, Strupp M, et al. Imaging human supraspinal locomotor centers in brainstem and cerebellum. Neuroimage. 2008;39: 786–792. doi:10.1016/J.NEUROIMAGE.2007.09.047

29. Spalek K, Coynel D, Freytag V, Hartmann F, Heck A, Milnik A, et al. A common NTRK2 variant is associated with emotional arousal and brain white-matter integrity in healthy young subjects. Transl Psychiatry. 2016;6: e758. doi:10.1038/TP.2016.20

30. Ohira K, Kumanogoh H, Sahara Y, Homma KJ, Hirai H, Nakamura S, et al. A truncated tropomyosin-related kinase B receptor, T1, regulates glial cell morphology via Rho GDP dissociation inhibitor 1. J Neurosci. 2005;25: 1343–1353. doi:10.1523/JNEUROSCI.4436-04.2005

31. Hamdan FF, Myers CT, Cossette P, Lemay P, Spiegelman D, Laporte AD, et al. High Rate of Recurrent De Novo Mutations in Developmental and Epileptic Encephalopathies. Am J Hum Genet. 2017;101: 664–685. doi:10.1016/j.ajhg.2017.09.008

32. Ogura H, Matsumoto M, Mikoshiba K. Motor discoordination in mutant mice heterozygous for the type 1 inositol 1,4,5-trisphosphate receptor. Behav Brain Res. 2001;122: 215–219. doi:10.1016/S0166-4328(01)00187-5

33. Van De Leemput J, Chandran J, Knight MA, Holtzclaw LA, Scholz S, Cookson MR, et al. Deletion at ITPR1 underlies ataxia in mice and spinocerebellar ataxia 15 in humans. PLoS Genet. 2007;3: 1076–1082. doi:10.1371/JOURNAL.PGEN.0030108

34. Krishnan A, Zhang R, Yao V, Theesfeld CL, Wong AK, Tadych A, et al. Genome-wide prediction and functional characterization of the genetic basis of autism spectrum disorder.Nat Neurosci. 2016;19: 1454–1462. doi:10.1038/nn.4353

35. Benton ML, Abraham A, LaBella AL, Abbot P, Rokas A, Capra JA. The influence of evolutionary history on human health and disease. Nature Reviews Genetics. Nature Research; 2021. p. 1. doi:10.1038/s41576-020-00305-9

36. Herrero J, Muffato M, Beal K, Fitzgerald S, Gordon L, Pignatelli M, et al. Ensembl comparative genomics resources. Database J Biol Databases Curation. 2016;2016. doi:10.1093/DATABASE/BAV096

37. Kent WJ, Sugnet CW, Furey TS, Roskin KM, Pringle TH, Zahler AM, et al. The Human Genome Browser at UCSC. Genome Res. 2002;12: 996–1006. doi:10.1101/GR.229102

38. Miga KH, Newton Y, Jain M, Altemose N, Willard HF, Kent EJ. Centromere reference models for human chromosomes X and Y satellite arrays. Genome Res. 2014;24: 697–707. doi:10.1101/GR.159624.113

39. Ernst J, Kellis M. ChromHMM: Automating chromatin-state discovery and characterization. Nature Methods. 2012. pp. 215–216. doi:10.1038/nmeth.1906

40. Quinlan AR, Hall IM. BEDTools: a flexible suite of utilities for comparing genomic features. Bioinformatics. 2010;26: 841–842. doi:10.1093/BIOINFORMATICS/BTQ033

41. Collins RL, Brand H, Karczewski KJ, Zhao X, Alföldi J, Francioli LC, et al. A structural variation reference for medical and population genetics. Nature. 2020;581: 444–451. doi:10.1038/s41586-020-2287-8

42. Yu G, Wang L-G, Han Y, He Q-Y. clusterProfiler: an R Package for Comparing Biological Themes Among Gene Clusters. Omi A J Integr Biol. 2012;16: 284–287. doi:10.1089/omi.2011.0118

43. He X, Sanders SJ, Liu L, De Rubeis S, Lim ET, Sutcliffe JS, et al. Integrated Model of De Novo and Inherited Genetic Variants Yields Greater Power to Identify Risk Genes. Williams SM, editor. PLoS Genet. 2013;9: e1003671. doi:10.1371/journal.pgen.1003671

44. Gokhman D, Kelman G, Amartely A, Gershon G, Tsur S, Carmel L. Gene ORGANizer: Linking genes to the organs they affect. Nucleic Acids Res. 2017;45: W138–W145. doi:10.1093/nar/gkx302

45. Song W, Lin GN, Yu S, Zhao M. Genome-wide identification of the shared genetic basis of cannabis and cigarette smoking and schizophrenia implicates NCAM1 and neuronal abnormality. Psychiatry Res. 2022;310. doi:10.1016/j.psychres.2022.114453

46. Finucane HK, Reshef YA, Anttila V, Slowikowski K, Gusev A, Byrnes A, et al. Heritability enrichment of specifically expressed genes identifies disease-relevant tissues and cell types. Nat Genet. 2018;50: 621–629. doi:10.1038/s41588-018-0081-4

47. Altshuler DM, Gibbs RA, Peltonen L, Schaffner SF, Yu F, Dermitzakis E, et al. Integrating common and rare genetic variation in diverse human populations. Nature. 2010;467: 52–58. doi:10.1038/nature09298

48. Weiner DJ, Gazal S, Robinson EB, O’Connor LJ. Partitioning gene-mediated disease heritability without eQTLs. Am J Hum Genet. 2022;109: 405–416. doi:10.1016/J.AJHG.2022.01.010

49. Li H, Handsaker B, Wysoker A, Fennell T, Ruan J, Homer N, et al. The Sequence Alignment/Map format and SAMtools. Bioinformatics. 2009;25: 2078–2079. doi:10.1093/bioinformatics/btp352

50. Reshef YA, Finucane HK, Kelley DR, Gusev A, Kotliar D, Ulirsch JC, et al. Detecting genome-wide directional effects of transcription factor binding on polygenic disease risk. Nat Genet. 2018;50: 1483. doi:10.1038/S41588-018-0196-7

